# Distinct adrenal gland macrophages regulate corticosteroid production

**DOI:** 10.1101/2023.09.05.556330

**Authors:** Conan JO O’Brien, Davi Sidarta-Oliveira, Aleksander Stawiarski, Giorgio Ratti, Pierre-Louis Loyher, Mélanie Makhlouf, Eman H AbouMoussa, Emma Haberman, Charles Sweeney, Yusuke Mizuno, Tomohiro Ishii, Tomonobu Hasegawa, Elise P Gomez-Sanchez, Celso E Gomez-Sanchez, Siamon Gordon, David R Greaves, Luis R Saraiva, Frederic Geissmann, Ana I Domingos

## Abstract

The adrenal glands are hormone secreting glands that sit on top of the kidneys. Adrenal glands produce glucocorticoids, mineralocorticoids, and catecholamines, and are therefore critical regulators of the stress response, the immune response, metabolism, and blood pressure. Despite being identified for more that 30 years, our understanding of adrenal macrophages remains incomplete. In numerous other tissues, macrophages carry out a plethora of physiological and homeostatic roles in addition to their classical immune functions. The aim of this study was to characterise the macrophage compartment of the adrenal gland and assess its contribution to adrenal function. Using an *in vivo* approach, we herein describe two morphologically and spatially distinct subsets of adrenal macrophages – dendritic-like macrophages that are present throughout the gland in young and old mice, and “foamy” lipid-laden macrophages that accumulate in the murine adrenal cortex in an age and diet-dependent manner. Furthermore, we present data showing that these foamy-like macrophages accumulate cholesterol and thereby regulate adrenal hormonal output, at steady state and in the context of obesity. We hereby provide novel insights into the physiological roles of macrophages in the adrenal gland and the mechanisms by which adrenal hormone production is regulated.

## Introduction

The adrenal glands are multi-hormone secreting glands situated superior to the kidneys. The glands are divided into the outer adrenal cortex and inner adrenal medulla which synthesise steroid hormones and catecholamines, respectively^1,2^. In mice and humans, the adrenal cortex, comprising the zona glomerulosa, the zona fasciculata, and zona reticularis (in humans only), serves as the sole producer of mineralocorticoids (aldosterone), and the primary producer of glucocorticoids - corticosterone in mice, cortisol in humans ^3,4^. The glucocorticoids are known regulators of the immune response, metabolism, and cognition, while aldosterone regulates blood pressure via controlling salt balance and total blood volume ^5,6^. The glucocorticoids are fundamentally regulated by the hypothalamic-pituitary-adrenal axis, and immediately controlled by adrenocorticotropic hormone (ACTH) ^7–11^, while aldosterone is regulated by the renin-angiotensin-aldosterone (RAAS) system, and immediately controlled by renin ^12–15^. It is well known that obesity causes alterations in corticosteroid responses which are thought to underly obesity-associated comorbidities such as hypertension and immune dysfunction ^16,17^.

Collectively, steroid hormones derived from the adrenal cortex are termed corticosteroids and all derive from cholesterol as the common precursor molecule. Cholesterol is metabolised into the corticosteroids intra-adrenally via a series of enzymatic reactions and its intra-adrenal levels are strongly associated with adrenal corticosteroid output ^18–20^. Cholesterol itself is a limiting factor in corticosteroid biosynthesis and its supplementation boosts corticosteroid production from adrenocortical cells *in vitro* ^21–23^. Currently, adrenal corticosteroid biosynthesis is believed to rely upon three cholesterol sources – intra-endocrine cell *de novo* synthesis of cholesterol, endocrine cell cholesterol stored in lipid droplets, and plasma lipoprotein-derived cholesterol ^18,24^.

Macrophages, though commonly known as the prototypical immune cell, are much more than “just” immune cells. In addition to their classical immune functions, macrophages carry out a range of physiological and homeostatic functions, complementary to the tissue in which they are found. Intestinal muscularis macrophages regulate peristalsis at steady state through secreted bone morphogenetic protein 2 ^25^. Embryonic-derived microglia directly and dynamically sculpt synapses and axonal outgrowth, pre- and postnatally ^26,27^. Similarly, embryonic-derived osteoclasts (bone macrophages) support bone development, resorption, and maintenance ^28^. In contrast to inflammatory recruited adipose tissue macrophages, adipose tissue-resident macrophages, through secreted platelet-derived growth factor c, directly promote lipid storage in mouse adipocytes, thereby regulating organismal body weight ^29^. Sympathetic neuron-associated macrophages in adipose tissues suppress lipolysis via the degradation of norepinephrine ^30^. Kupffer cells (liver-resident macrophages), via scavenger receptors, and CCR2-dependent recruited liver macrophages, remove dead and damaged erythrocytes and their components from the circulation and recycle iron into hepatocytes ^31,32^.

Macrophages were first identified in the adrenal medulla and cortex of both mice and humans in 1984 and 1994, respectively ^33,34^. They describe macrophages that fall into one of two general categories: those located adjacent to vascular sinuses and those located throughout the parenchyma. The parenchymal macrophages extend dendrites though, and come into direct contact with parenchymal cells ^33,34^, indicating potential paracrine interactions. However, to date, few studies have directly characterised and probed the function of the macrophage compartment of the murine adrenal gland.

Here, we present data describing two morphologically, spatially, and phenotypically distinct adrenal macrophage populations that reside in the adrenal cortex. CX3CR1-expressing Zona Glomerulosa Macrophages (ZGMs) express corticosteroid biosynthetic enzymes and we show that their depletion blunts aldosterone levels at steady state in adolescent (10-week-old) mice. Big Autofluorescent Lipid-Laden (BALL) macrophages accumulate in the adrenal cortex in an age-dependent manner and contain cholesterol, the necessary precursor molecule of the corticosteroids. Using salt and angiotensin II-mediated modulations of aldosterone production we show that BALL macrophages dynamically alter their size and cholesterol contents to meet the demands of endocrine cells. CX3CR1-mediated ablation of the BALL macrophage compartment, but not the ZGM compartment, cuts intra-adrenal cholesterol stores by half and suppresses corticosteroid levels in adult mice (20 week plus) at steady state and in obesity. Thus, adrenal macrophages prove to be unforeseen novel players in the HPA and RAAS systems with potential therapeutic relevance in endocrine disorders.

## Results

### The adrenal cortex harbours two morphologically distinct macrophage subsets

Our initial aim was to characterise the macrophage compartment of the murine adrenal gland morphologically and spatially. We carried out immunofluorescent staining and confocal microscopy in the adrenal glands of mice expressing GFP from CX3CR1-expressing cells, a widely used mouse line that labels microglia and tissue macrophages ^35^. CX3CR1-expressing macrophages were located throughout the adrenal gland, with particularly high densities observed in the zona glomerulosa and the medulla (Fig. 1A). Staining against the pan-macrophage marker CD68 revealed an additional macrophage subset revealed an additional, non-overlapping subset that preferentially localised to the zona fasciculata and to a lesser extent the zona glomerulosa (Fig. 1B). CX3CR1-expressing zona glomerulosa macrophages (ZGMs) have the classical dendritic morphology and come into direct contact with endocrine cells expressing CYP11B2 (Fig. 1B), alternatively known as aldosterone synthase, indicating a potential paracrine interaction between these two cell types. Flow cytometric analysis showed that CX3CR1+ ZGMs remain constant in number and as a frequency of the total CD68+ macrophage compartment at 10 and 15 weeks of age, with a slight decrease observed at 20 weeks of age (Fig. 1C). We observed an age-dependent accumulation of CD68^++^ macrophages in the zona fasciculata, with cells becoming more populous and bigger in size with age (Fig. 1D). Flow cytometry revealed that CX3CR1-negative macrophages were significantly increased in number and proportion in the adrenal cortexes of 20-week-old mice (Fig. 1E). In contrast to ZGMs, CD68^++^ cortical macrophages have a “foamy” morphology, similar to the cholesterol-loaded foam cell macrophages that characterise atherosclerotic plaques. These macrophages have a highly autofluorescent cellular content, which is located in intracellular vacuoles (Fig. 1F). These two phenotypically and morphologically distinct macrophage populations co-reside in the adrenal cortex (Fig. 1G). Additionally, they differ on the basis of Ki-67 expression, with CX3CR1+ ZGMs being Ki-67^lo^ and CX3CR1-negative BALL macrophages being Ki-67^hi^ at two out of the three timepoints tested (Fig. S1A).

**Figure 1.**
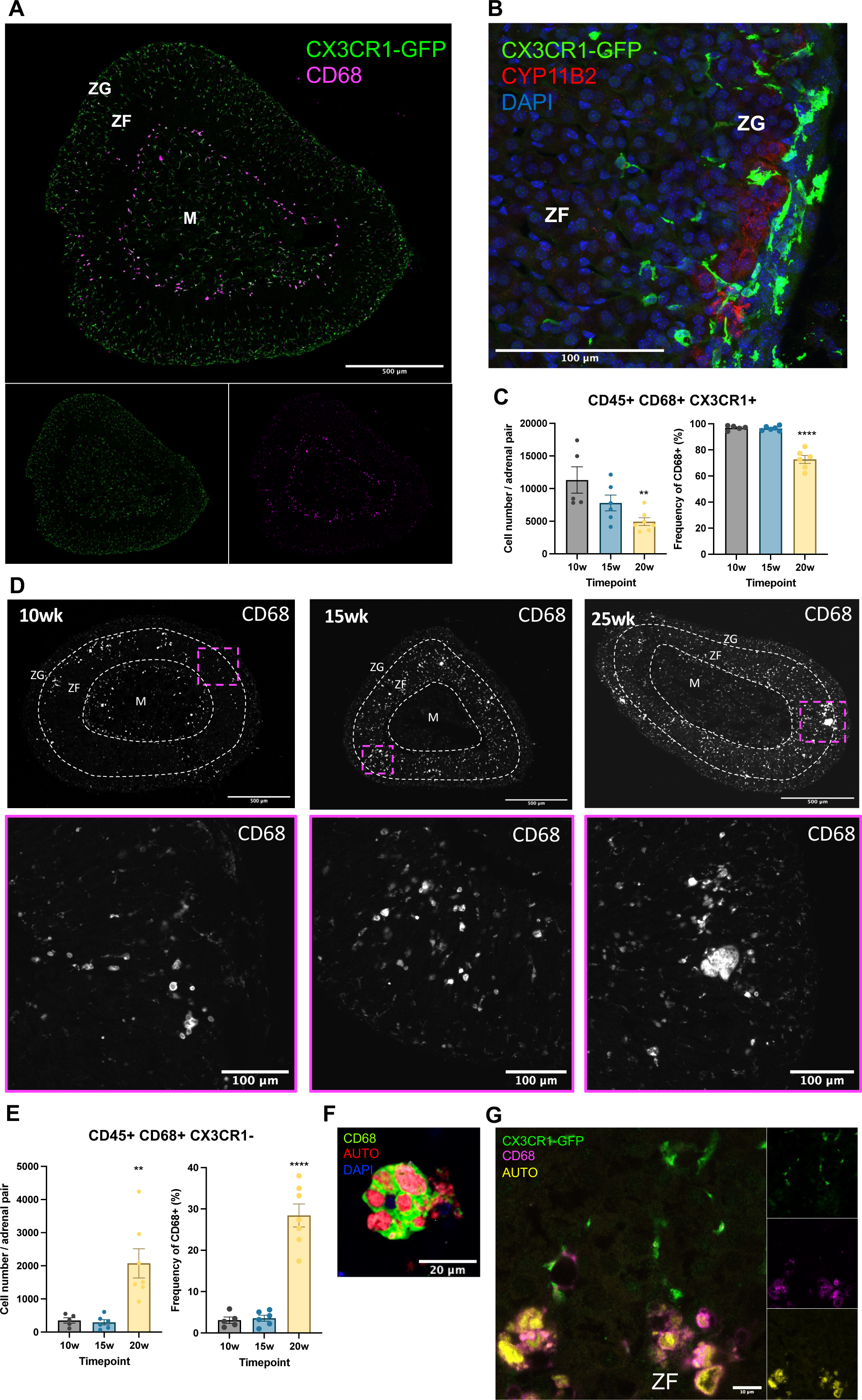
The adrenal cortex harbours two morphologically distinct macrophage subsets. (A) A representative confocal image showing a cross section of a mouse adrenal gland isolated from a 12 week old CX3CR1^GFP^ mouse and stained using an anti-GFP (green) and an anti-CD68 (white) antibody. (B) A representative confocal image focusing on the adrenal zona glomerulosa of a CX3CR1^GFP^ mouse and stained with an anti-GFP (green) antibody, an anti-CYP11B2 (red) antibody, and DAPI (blue). Scale bars in (A) and (B) are 100μm. (C) Flow cytometry data showing the absolute number of CD45+ CD68+ CX3CR1+ cells (left), and frequency of CX3CR1+ cells among CD45+ CD68+ cells (right) in the adrenal cortex at the indicated timepoints. Each data point depicts the data from one mouse. (D) Representative confocal images showing cross sections of wild-type mouse adrenal glands at the indicated timepoints stained against CD68 (white). White dotted lines indicate demarcation of the different zones of the adrenal gland. Dashed magenta boxes in the above images for each timepoint indicate the areas that are enhanced in the magenta boxes below. Scale bars in the top images are 500μm and the bars in the below images are 100μm. (E) Flow cytometry data showing the absolute number of CD68+ CX3CR1-cells (left) and frequency of CX3CR1-cells among CD45+ CD68+ cells (right) in the adrenal cortex at the indicated timepoints. Each data point depicts the data from one mouse. (F) A close-up confocal image of a cortical CD68+ (green) macrophage showing the autofluorescent (red) contents in intracellular vacuoles. DAPI is shown in blue – scale bar 20μm. (G) A representative confocal image of the adrenal cortex of a 22 week old male CX3CR1^GFP^ reporter mouse stained with anti-GFP (green), anti-CD68 (magenta) and autofluorescence shown in yellow - scale bar 10μm. Data in C and E were analysed by one way ANOVA with Tukey’s test for multiple comparisons and are shown as average ± s.e.m. \**P* < 0.05, \*\**P* < 0.01, \*\*\**P* < 0.001, \*\*\*\**P* < 0.0001. ZG = zona glomerulosa; ZF = zona fasciculata; M = medulla; AUTO = autofluorescence.

The adrenal gland is widely known to be autofluorescent, which is attributed to its high lipid content ^36^. To ensure that the autofluorescent macrophages were not an artefact of fluorescent imaging, we characterised the autofluorescent signature of the aged (20-week-old) adrenal gland, finding wide spectral autofluorescent signals which decline as we approach the far-red end of the light spectrum (Fig. S1B). By staining with a directly conjugated anti-CD68 antibody with a fluorophore that emits outside the adrenal autofluorescent signal, we confirmed that the adrenocortical macrophages were not an artefact and were the source of the autofluorescent signal inherent to the aged adrenal gland (Fig. S1C). We additionally verified CD68 staining in the adrenal gland using classical immunohistochemistry with Fast Red staining (Fig. S1D). Taken collectively, the data suggest that the adrenal cortex harbours two distinct macrophage populations – dendritic <u>Z</u>ona <u>G</u>lomerulosa <u>M</u>acrophages (ZGMs) and <u>B</u>ig <u>A</u>utofluorescent <u>L</u>ipid-<u>L</u>aden (BALL) macrophages that resemble cholesterol-containing foamy macrophages, that can be differentiated by CX3CR1 expression and their autofluorescent signature.

### CX3CR1-expressing progenitors seed long lived BALL macrophages and ZGMs

Our next objective was to establish the ontogeny of the priorly identified macrophage populations. First, using a single cell RNA seq dataset comprising CD45+ leukocytes in the adrenal gland ^37^ we used *Lyz2*, *Ccr2*, *Cd68* and *Cx3cr1* as myeloid markers and an unsupervised cell-type annotation tool ^38^ to filter 6,377 myeloid cells for downstream analyses. Further sub-clustering of these myeloid cells identified three distinct populations of myeloid cells in the adrenal gland (Fig. S2A). All three populations expressed *Cd68,* while populations one and two express *Cx3cr1* (Fig. 2A). Using RNA velocity, we performed trajectory analysis on this myeloid cell fraction, which suggests that populations two and three derive from population one, a Cx3cr1+ population that expresses the mature macrophage markers *Mrc1, Cd36*, and *Cd68* (Fig. 2B). Next, we sought to establish experimentally whether the CX3CR1+ ZGMs and BALL macrophages can be replaced after irradiation-induced injury and redistribute to the same spatial location within the adrenal. We performed an irradiation and bone marrow transplantation experiment in which 9-week-old wild-type CD45.1-expressing mice were irradiated, then injected with bone marrow from CD45.2-expressing CD68^GFP^ reporter mice ^39^. Then, 11 weeks later, we analysed the adrenal macrophage compartment by immunofluorescent imaging and flow cytometry (Fig. 2C). We examined the (F4/80+ MHCII+) macrophage compartment in the adrenal gland, kidney, liver, and spleen, finding similar levels of chimerism across all tissues (Fig. 2D). By imaging, we observed that BALL macrophages were largely GFP+ (Fig. 2E), indicating that they derived from the transplanted bone marrow. We also observed repopulation of the ZGM compartment. CX3CR1-positive and negative adrenal macrophages differed in their frequency chimerism, with CX3CR1-negative macrophages (BALLs) deriving from the bone marrow at a significantly higher rate than CX3CR1+ macrophages (ZGMs) (Fig. 2F). To test the hypothesis that the CX3CR1-negative BALL macrophages develop from a CX3CR1-positive precursor, as predicted by our RNA velocity trajectory analysis (Fig. 2B), we performed a linage tracing experiment. Five- to six-month-old Cx3cr1-CreERT; Rosa26mTmG mice were pulsed with tamoxifen to induce recombination in CX3CR1-expressing cells, which tags recombined cells and their progeny with GFP. Two months later, adrenals were harvested. We observed GFP from macrophages in the zona glomerulosa, as well as autofluorescent cells in the zona fasciculata (Fig. 2H). Approximately 50% of total adrenal gland macrophages expressed GFP, similar to kidney macrophages, and at a slightly lower rate than microglia (Fig. 2I). To quantitatively assess whether BALL macrophages come from CX3CR1+ precursors, we measured the frequency of autofluorescent cells in each adrenal region. In agreement with our prior data, the vast majority of autofluorescent cells situated in the zona fasciculata (Fig. 2J). Recombination among autofluorescent cells occurred at ∼50% in all adrenal regions (Fig. 2K). Taken collectively, this data indicate that both ZGMs and BALL adrenal gland macrophages undergo repopulation from the bone marrow, albeit at slightly different rates. Adrenal gland macrophages as a whole are a long lived-cell type, similar to macrophages from other tissues. Additionally, the data indicate that BALL macrophages derive from a CX3CR1-expressing precursor cell.

**Figure 2.**
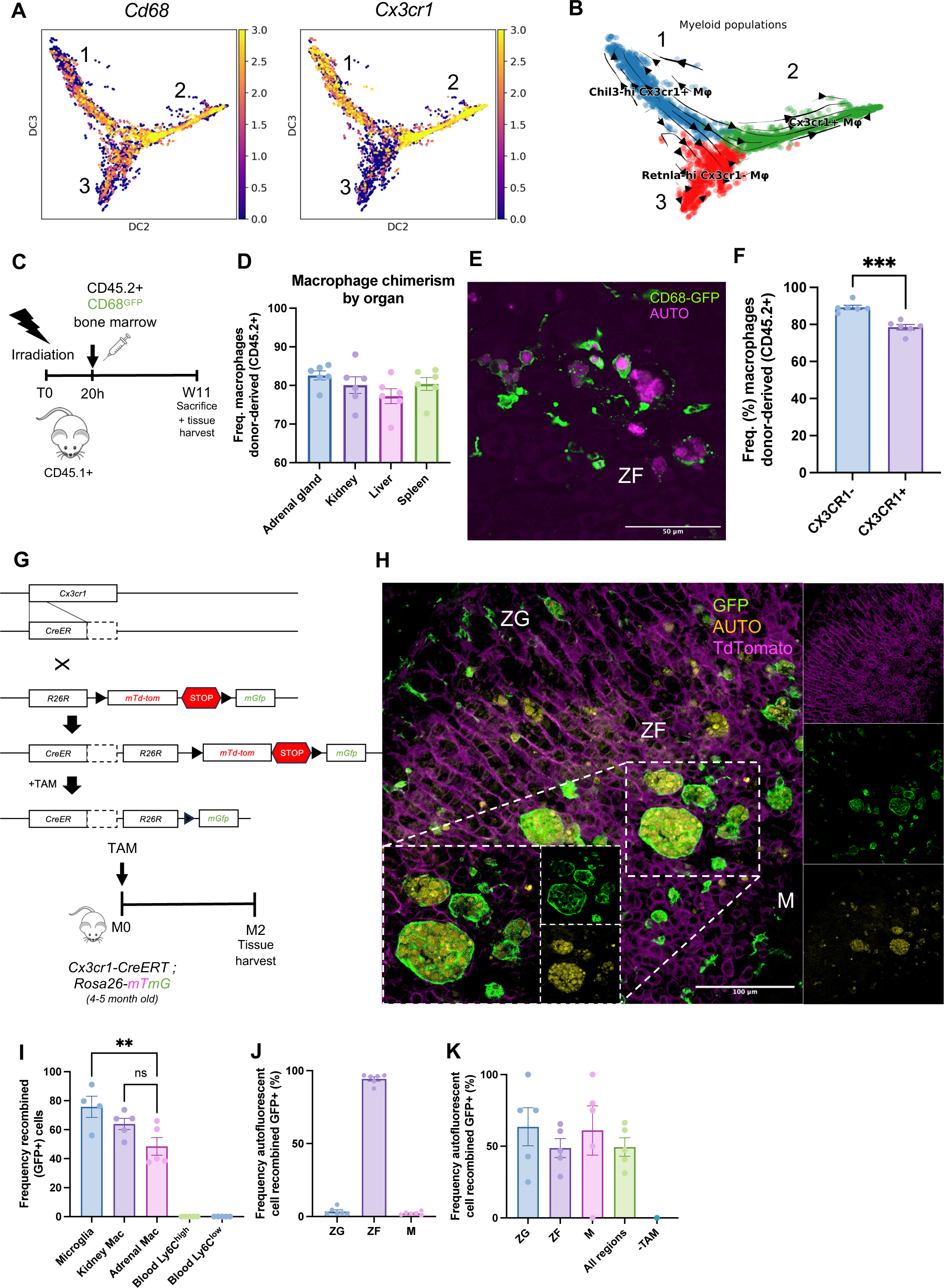
CX3CR1-expressing progenitors seed long lived BALL macrophages and ZGMs. (A) Heat maps showing the expression of Cd68 and Cx3cr1 genes in adrenal myeloid cells. (B) Single cell RNA seq plot showing RNA trajectories of the myeloid compartment of the adrenal gland. (C) A schematic depicting the experimental procedure followed for panels D-F. Wild-type mice were irradiated then rescued by injection with CD68-GFP bone marrow and culled 11 weeks later. (D) Bar chart showing CD45+ F4/80+ macrophage chimerism by organ. (E) Representative confocal image showing GFP+ autofluorescent macrophages in the adrenal cortex of mice that underwent bone marrow transplantation. (F) Bar chart with frequencies of CD45.2+ (donor-derived) cells in CX3CR1+ and CX3CR1-adrenal macrophages. (G) Schematic representation of the mouse cross used to generate the Cx3cr1-CreERT; Rosa26mTmG mice, and the experimental timeline used to generate the data in H-K. (H) Representative confocal image showing mouse adrenals from the experiment depicted in panel (G) with GFP (green) autofluorescence (yellow) and TdTomato (magenta). (I) Frequency of CD45+ F4/80+ CD64+ tissue macrophages or blood monocytes that are recombined (GFP+) 2 months following tamoxifen administration. (J) Frequency of autofluorescent cells divided by region in the adrenal gland. (K) Frequency of autofluorescent cells that have recombined (GFP+) in each adrenal region two months following tamoxifen administration. ZG = zona glomerulosa; ZF = zona fasciculata; M = medulla; AUTO = autofluorescence. Data in F were analysed by two-tailed unpaired Student’s *t*-test and are shown as average ± s.e.m. Data in I and K were analysed by one way ANOVA with Tukey’s test for multiple comparisons and are shown as average ± s.e.m. \**P* < 0.05, \*\**P* < 0.01, \*\*\**P* < 0.001, \*\*\*\**P* < 0.0001.

### CX3CR1+ Zona Glomerulosa Macrophages (ZGMs) regulate aldosterone production but not other adrenal-derived hormones

We next examined the gene expression patterns of CX3CR1-expressing ZGMs using FACS sorting and bulk RNA-seq. From the adrenal glands of CX3CR1^GFP/GFP^ mice we sorted CD45-cells (endocrine fraction) and CD45+ MHCII+ F4/80+ CX3CR1+ ZGMs (Fig. 3A). As expected, we detected abundant expression of the macrophage marker genes *H2-Ab1, Lyz2, Csf1r, Cd68,* and *Adgre1* in the sorted ZGMs, in contrast to the endocrine fraction (Fig. 3B).

**Figure 3.**
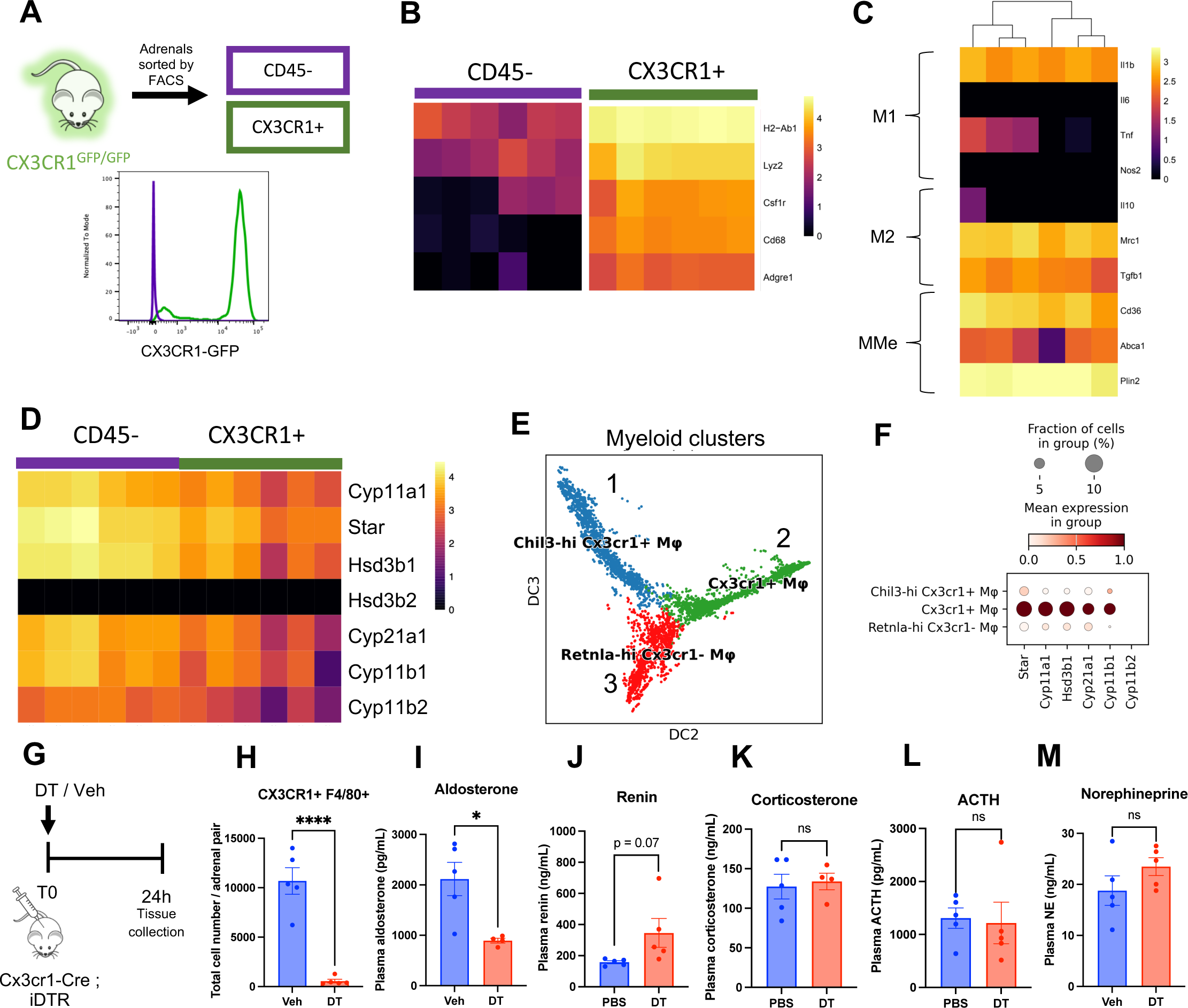
CX3CR1+ Zona Glomerulosa Macrophages (ZGMs) regulate aldosterone production, but not other adrenal-derived hormones. (A) A schematic (above) depicting the experimental outline for the data shown in figures B-D. CD45-(purple; endocrine) and CX3CR1+ (green) cells were sorted from the adrenal glands of CX3CR1^GFP/GFP^ mice for bulk RNA sequencing. The histogram below shows the CX3CR-GFP expression in the sorted cell populations. (B) Heatmaps showing normalized expression of the indicated macrophage marker genes (B), polarization marker genes (C) and corticosteroid biosynthetic genes (D) in the CD45- and CX3CR1+ cell populations. Each box in the horizontal axis represents data from one mouse. Expression levels are represented on a log10 scale of normalized counts (NC) plus one. (E) Single cell RNA seq data of the adrenal myeloid compartment. (F) Single cell RNA seq dot plots showing the standardised expression of adrenal corticosteroid biosynthetic genes in the indicated myeloid populations. (G) A schematic showing the experimental procedure followed for the data shown in H-M. Cx3cr1-Cre; iDTR mice were injected with diphtheria toxin (DT) or vehicle (Veh) control and tissues were collected from mice 24 hours later. (H) Flow cytometry data quantifying the total number of CX3CR1+ F4/80+ cells in Vehicle or DT-treated mice. Bar charts showing the levels of aldosterone (I), renin (J), corticosterone (K), ACTH (L), and norepinephrine (M) in the blood of vehicle or DT-treated mice. Each data point denotes data from one mouse. Data in H-M were analysed by two-tailed unpaired Student’s *t*-test and are shown as average ± s.e.m. \**P* < 0.05, \*\**P* < 0.01, \*\*\**P* < 0.001, \*\*\*\**P* < 0.0001.

We next interrogated ZGM gene expression in line with the M1/M2/metabolic activation paradigm ^40^. We found that ZGMs exhibit a mixed phenotype, characterised by expression of the M1 marker *Il1b,* the M2 markers *Mrc1* (CD206) and *Tgfb1,* and the metabolic activation markers *Cd36* and *Plin2* (Fig. 3C). As expected, we observed high expression of corticosteroid biosynthetic enzyme genes in the CD45-endocrine fraction, but to our surprise we detected substantial levels of these transcripts, most notably *Cyp11a1*, *Star*, and *Hsd3b1*, in Zona Glomerulosa macrophages (Fig. 3D). Next, using the single cell RNA-seq data of filtered myeloid cells (Fig. 2A, B, S2A), we examined the gene expression of corticosteroid synthesis genes among the three adrenal myeloid populations (Fig. 3E). In agreement with our bulk RNA-seq data, gene expression analysis showed that the Cx3cr1+ macrophages (population two) expressed relatively high levels of the corticosteroid biosynthetic genes, except for Cyp11b2 (Fig. 3F, S2B). Since the data to this point suggest that ZGMs express the machinery to contribute to the synthesis of adrenal corticosteroids, we next assessed the contribution of ZGMs to adrenal hormonal output. We crossed Cx3cr1-Cre mice with Cre-inducible diphtheria toxin receptor mice to generate Cx3cr1-Cre; iDTR mice. At 10 weeks we injected the mice with 750ng diphtheria toxin (DT) or vehicle control and sacrificed the mice 24 hours later (Fig. 3G). We validated depletion of ZGMs by flow cytometry, observing a significant and almost complete ablation of ZGMs in DT-treated mice relative to controls (Fig. 3H). In the plasma, we observed a significant reduction in aldosterone concentration (Fig. 3I), with the expected compensatory relative increase in renin in DT-treated mice (Fig. 3J). Corticosterone, ACTH, and norepinephrine levels were unaffected by ZGM depletion (Fig. 3K-M). Taken collectively, these findings indicate that ZGMs contribute to adrenal aldosterone output via the synthesis of steroid precursors to the endocrine cells of the zona glomerulosa.

### BALL macrophages contain cholesterol and their formation is accelerated in obesity

We previously established that Big Autofluorescent Lipid-Laden (BALL) macrophages accumulate in the adrenal cortex under steady state in an age-dependent manner (Fig. 1). We hypothesised that the lipid contained in BALL macrophages is cholesterol, given its metabolic and physiological relevance in the adrenal gland as the necessary precursor molecule to the adrenal steroid hormones corticosterone and aldosterone (Fig. 4A). A previous study ^41^ generated mice (Cre-ER^TM^; Star^flox/flox^mice) with a tamoxifen-induced knock-out of the steroidogenic acute regulatory protein (*Star*) gene in adult mice (Fig. 4A). They observed extensive lipid accumulation in large vacuolated cells in the adrenals of these mice, driven by accumulating cholesterol that can not be metabolised into the steroid hormones. We stained these adrenal glands against the pan macrophage marker CD68 and found that the accumulating lipid / cholesterol in these adrenals is contained entirely inside macrophages (Fig. 4B). This macrophage-bound cholesterol was highly autofluorescent (Fig. S3A), in agreement with our previous findings (Fig. S1A). Next, we wanted to determine the effect of increased plasma cholesterol levels in BALL macrophages. To induce hypercholesterolaemia we fed mice a high fat diet (HFD) for 12 weeks. HFD-fed mice had elevated total plasma cholesterol levels and plasma cholesterol esters, relative to normal diet-fed mice (Fig. S3B, S3C). HFD-fed mice also had accelerated BALL macrophage formation, and the cells were significantly more autofluorescent and lipid-laden than BALL macrophages in normal diet-fed mice (Fig. 4C-E).

**Figure 4.**
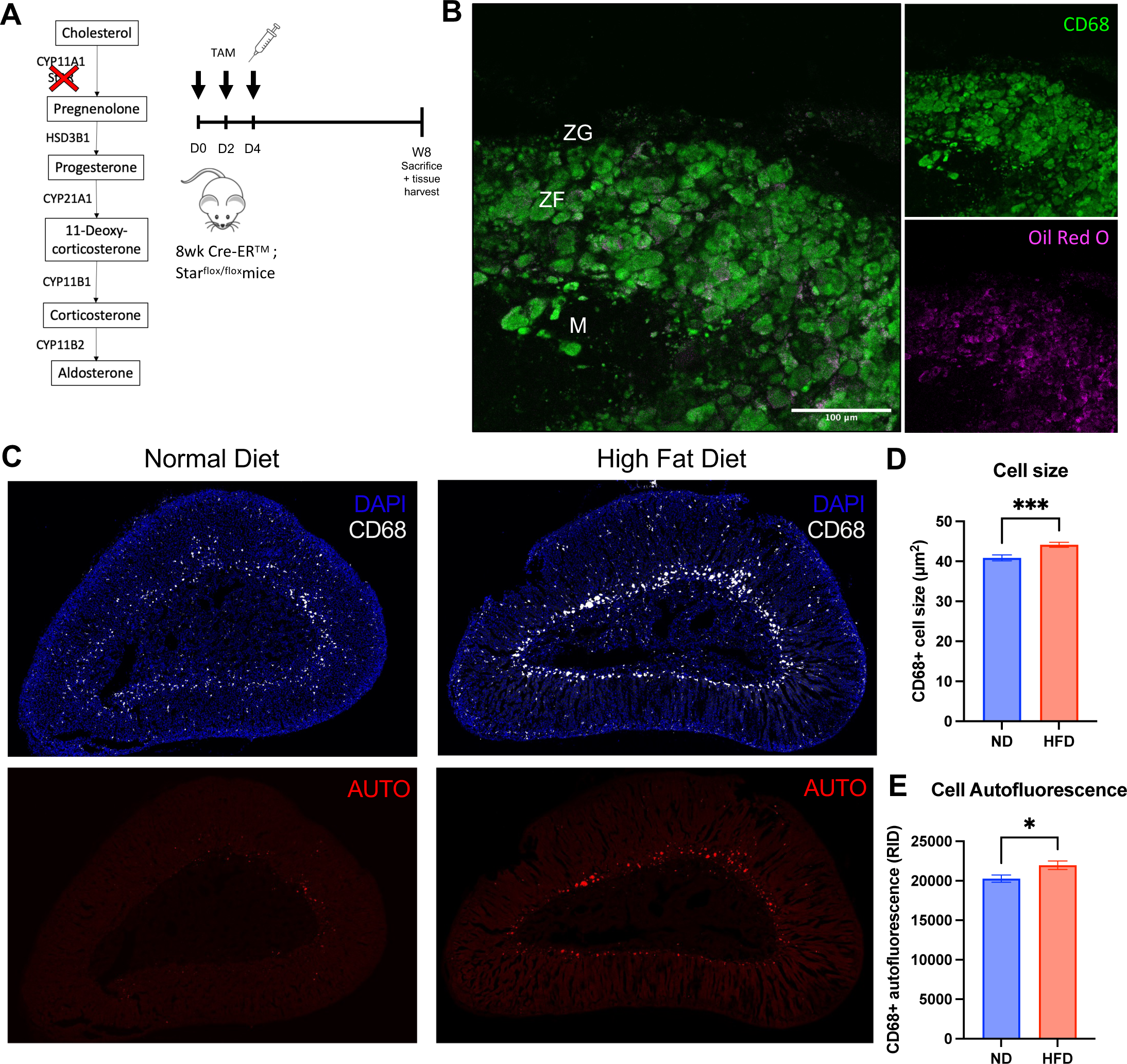
BALL macrophages contain cholesterol and their formation is accelerated in obesity. (A) Schematic depicting the experimental protocol used to generate the tissue that was analyzed in panel B. Eight week old CAGGCre-ER^TM^; Star^flox/flox^ mice were pulsed with tamoxifen on days 0, 2, and 4 to induce temporal deletion of Star, the rate-limiting enzyme in the adrenal corticosteroid biosynthetic pathway. Eight weeks later adrenal glands were harvested for subsequent microscopic analyses. (B) A representative confocal image showing an adrenal gland from a CAGGCre-ER^TM^; Star^flox/flox^ mouse stained with anti-CD68 (green) and Oil Red O (magenta). (C) Representative confocal images of adrenal cross sections from 22-week-old male CX3CR1^GFP^ mice that were fed a normal diet (ND) or high fat diet (HFD) for 12 weeks, with DAPI (blue), anti-CD68 staining (white) and autofluorescence (red). Bar charts quantifying the mean CD68+ cell size (D) and autofluorescence (E) in ND and HFD-fed mice. Data in D and E were analysed by two-tailed unpaired Student’s *t*-test and are shown as average ± s.e.m. \**P* < 0.05, \*\**P* < 0.01, \*\*\**P* < 0.001, \*\*\*\**P* < 0.0001.

### Corticosteroid stimulation and inhibition differentially impact BALL macrophage size and cholesterol contents

We next probed how corticosteroid stimulation and inhibition impacted BALL macrophage size and cholesterol contents. To suppress corticosteroid production, we fed CX3CR1^GFP/+^ mice a high salt diet (HSD) for six days (Fig. 5A). HSD-fed mice had significantly reduced levels of plasma aldosterone (Fig. 5C) and marginally reduced plasma corticosterone relative to normal salt diet-fed mice (Fig. 5D). HSD-fed mice had significantly larger and more autofluorescent BALL macrophages relative to NSD-fed mice (Fig. 5E, F).

**Figure 5.**
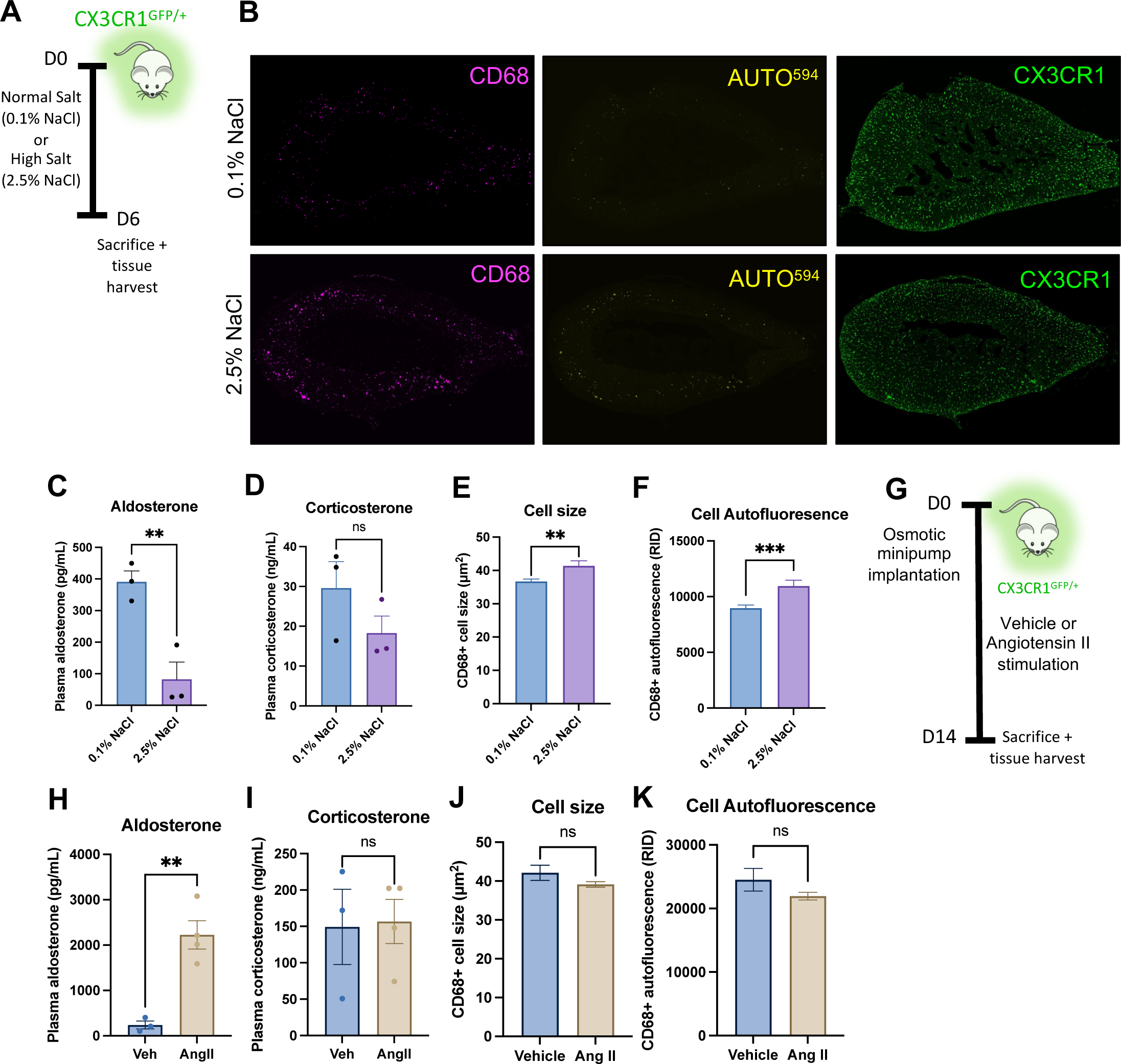
Corticosteroid stimulation and inhibition differentially impact BALL macrophage size and cholesterol contents. (A) Schematic depicting the experimental protocol used to generate the tissue that was analyzed in panel B-F. CX3CR1^GFP/+^ mice were fed either a normal (0.1% NaCl) or high salt (2.5% NaCl) diet for six days before tissue harvest for subsequent analyses. (B) Representative confocal images of adrenal glands from the indicated experimental groups with anti-CD68 staining (magenta), autofluorescence (yellow), and anti-GFP staining (green). Bar charts showing plasma aldosterone (C) and corticosterone (D) levels in the indicated treatment groups. Bar charts showing the average cell size (E) and average cell autofluorescence (K) in the indicated treatment groups. (G) On day 0 mice were implanted subcutaneously with an osmotic minipump containing either vehicle or Ile^5^– Angiotensin II (AngII; 1000 ng/kg per minute) for 14 days, prior to sacrifice and tissue harvest. Bar charts showing plasma aldosterone (H) and corticosterone (I) levels in the indicated treatment groups. Bar charts showing the mean cell size (J) and mean cell autofluorescence (K) in the indicated treatment groups. Each data point denotes data from one mouse. Data in C-F and H-K were analysed by two-tailed unpaired Student’s *t*-test and are shown as average ± s.e.m. \**P* < 0.05, \*\**P* < 0.01, \*\*\**P* < 0.001, \*\*\*\**P* < 0.0001.

To stimulate aldosterone production, we subcutaneously implanted an osmotic minipump containing Ile^5^–Angiotensin II (AngII; 1000 ng/kg per minute) or vehicle control (Fig. 5G) ^42^. AngII-treated mice had significantly elevated plasma aldosterone concentrations relative to controls, while corticosterone levels remained unchanged (Fig. 5H, I). BALL macrophages in AngII-treated mice had trends towards reductions in BALL cell size and autofluorescence. Taken collectively, these findings indicate that BALL macrophages dynamically respond to corticosteroid stimulation and inhibition, with changes apparent in BALL macrophage size and cholesterol contents.

### CX3CR1 regulates BALL formation, intra-adrenal cholesterol stores, and corticosteroid output

In light of the observation that BALL macrophages can dynamically respond to adrenal corticosteroid output, we next tested the hypothesis that BALL macrophages support adrenal hormonal output via the provision of cholesterol. While the CX3CR1-GFP line is commonly used as a macrophage reporter, as a knock-in-knock-out mouse line ^35^ mice homozygous for the transgene are functional knockouts for the *Cx3cr1* gene. We observed that CX3CR1^GFP/GFP^ mice had almost an entire ablation of BALL macrophages in the adrenal cortex relative to wild type mice (Fig. 6A). When BALL macrophages were present, they were significantly smaller in size and in autofluorescent cholesterol contents when compared to BALLs in wild-type mice (Fig 6B, C). To test whether CX3CR1-deficient macrophages were inherently defective in their ability to uptake cholesterol, we treated WT or CX3CR1^GFP/GFP^ bone marrow-derived macrophages with fluorescent BODIPY-FL Low Density Lipoprotein and assessed their ability to take up cholesterol. We found no difference between the two genotypes in their ability to form take up cholesterol and form foamy-like cells *in vitro* (Fig. S4A). Adrenal corticosteroid biosynthetic enzyme expression was mostly unchanged between wild type and CX3CR1^GFP/GFP^ mice, except for *Hsd3b1* which was significantly elevated in knockout mice relative to controls (Fig. 6D). While adrenal weights did not differ between wild type and knockout mice (Fig. 6E), CX3CR1^GFP/GFP^ mice had intra-adrenal cholesterol stores approximately half the levels of wild-type controls (Fig. 6F), consistent with the ablation of cholesterol-containing BALL macrophages. Systemic plasma cholesterol titres were similar between the genotypes (Fig. 6G). Consistent with reduced intra-adrenal cholesterol stores, CX3CR1^GFP/GFP^ mice had significantly reduced plasma aldosterone and corticosterone concentrations relative to wild-type mice at steady state (Fig. 6H, I). As expected, we observed a compensatory increase in plasma renin in knockout animals relative to wild-type (Fig. 6J) mice, with ACTH levels unchanged between the two genotypes (Fig. 6K)

**Figure 6.**
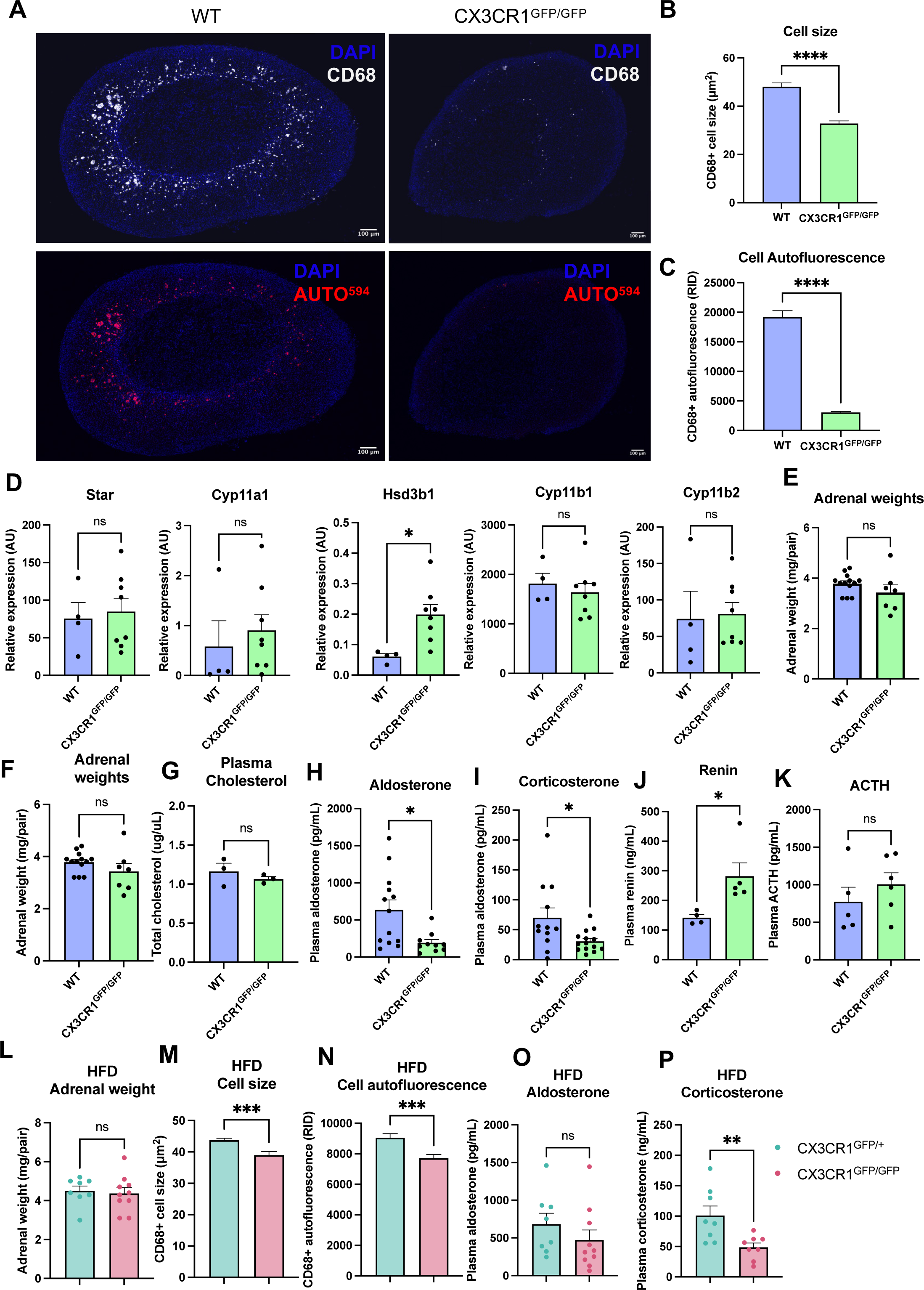
CX3CR1 regulates intra-adrenal cholesterol stores, BALL formation, corticosteroid output. Representative confocal images of adrenal cross sections from 22-week-old male wild type and CX3CR1^GFP/GFP^ mice that were fed a normal with DAPI (blue), anti-CD68 staining (white)m and autofluorescence (red). Bar charts quantifying the mean CD68+ cell size (B) and autofluorescence (C) in the indicated genotypes. (D) RT-qPCR data showing the relative expression of the indicated corticosteroid biosynthetic genes. Bar charts showing the adrenal weights (E), intra-adrenal total cholesterol levels (F), plasma cholesterol levels (G), plasma aldosterone (H), corticosterone (I), renin (J), and ACTH (L) levels in the indicated genotypes. Bar charts showing the adrenal weights (M), mean CD68+ cell size (N) and autofluoresence (O), plasma aldosterone concentrations (P) and plasma corticosterone concentrations (Q) in high fat diet-fed mice of the indicated genotypes. Each data point denotes data from one mouse. All pairwise comparisons were analysed by two-tailed unpaired Student’s *t*-test and are shown as average ± s.e.m. \**P* < 0.05, \*\**P* < 0.01, \*\*\**P* < 0.001, \*\*\*\**P* < 0.0001.

Finally, we put CX3CR1^GFP/+^ and CX3CR1^GFP/GFP^ mice on a high fat diet for 12 weeks to test whether BALL macrophages regulate corticosteroid responses in the context of obesity. CX3CR1^GFP/GFP^ had marginally higher total body weights compared to CX3CR1^GFP/+^ mice (Fig. S4b). Like in steady state, adrenal weights did not differ between CX3CR1-sufficient and CX3CR1-deficient animals (Fig. 6L), but BALL cell size and autofluorescent cholesterol contents were significantly reduced in knockouts relative to control mice (Fig. 6M, N). Plasma aldosterone concentrations did not differ between the two genotypes, but corticosterone concentrations were significantly reduced in CX3CR1 knockout mice relative to controls (Fig. 6O, P). Together, these results indicate that CX3CR1 regulates the formation of BALL macrophages and the establishment of additional intra-adrenal cholesterol stores. Moreover, these stores appear necessary for supporting adrenal corticosteroid output at steady state and in obesity.

## Discussion

Our observations in the present study indicate that distinct populations of adrenal gland macrophages regulate adrenal hormonal output in steady state and under the pathological state of obesity. CX3CR1+ Zona Glomerulosa Macrophages (ZGMs), the predominant macrophage subset in adolescent (10-week-old) mouse adrenals, express corticosteroid biosynthetic genes and contribute towards aldosterone output at steady state. Morphologically distinct Big Autofluorescent Lipid-Laden (BALL) macrophages descend from a CX3CR1-expressing precursor, accumulate in the adrenal cortex in an age and diet-dependent manner, and regulate corticosteroid output at steady state and in obesity in adult mice (20 weeks plus).

We show that ZGMs positively regulate aldosterone production at steady state via expression of corticosteroid biosynthetic enzymes, however according to our data ZGMs themselves do not express the last enzyme in the aldosterone biosynthetic pathway, aldosterone synthase (*Cyp11b2*). Our data therefore suggest that ZGMs provide the precursors to aldosterone and in this way support aldosterone production. CYP11B1 catalyses the synthesis of corticosterone, which in turn can be converted to aldosterone by CYP11B2. In this way, aldosterone production via CYP11B1 is a legitimate biosynthetic route, as evidenced by *Cyp11b1* deletion in mice significantly suppressing aldosterone production ^43^. CX3CR1 and biosynthetic enzyme-expressing macrophages preferentially localise to the zona glomerulosa, which may explain why the depletion of CX3CR1-expressing macrophages did not impact corticosterone levels in our study. Macrophages have previously been linked with playing a role in the renin-angiotensin-aldosterone-system (RAAS). Op/Op mice lack the colony stimulating factor 1 gene (*Csf1*), a key survival factor for tissue macrophages, are resistant to Angiotensin II-induced elevations in blood pressure, in contrast to wild type mice ^42^. This is consistent with our findings that ZGMs situate adjacent to the aldosterone producing cells of the zona glomerulosa and that they express corticosteroid biosynthetic genes. This may therefore offer a new therapeutic angle for treating hypertension caused by elevated aldosterone or hyperaldosteronism.

While a link between circulating cholesterol levels and adrenal corticosteroid output may seem intuitive, linking the two is not straightforward. According to our data, BALL macrophages comprise a sizeable intra-adrenal cholesterol store that is distinct from endocrine cell cholesterol stores and the circulating pool of cholesterol found in plasma. We show that this store dynamically changes in size with differing corticosteroid output demands. Further, our data indicate that the reductions in corticosteroid output observable in BALL macrophage-deficient adrenals are attributable to diminished intra-adrenal cholesterol stores, rather than fluctuations in renin, ACTH, or corticosteroid biosynthetic enzymes, the major known regulators of corticosteroid output. CX3CR1 was previously shown to promote atherogenesis and the formation and survival of foam cells in atherosclerotic plaques ^44^. Here, we present evidence of another context in which CX3CR1 inhibition could be beneficial – via suppression of intra-adrenal cholesterol stores.

The concept of intra-adrenal cholesterol stores being directly linked to corticosteroid output is not a new one. A recent paper reported that Plin2, a gene that promotes formation of lipid droplets, negatively regulates intra-adrenal cholesterol and cholesterol ester stores, as well as corticosterone output ^20^. While this study focused on analysing the size of lipid droplets within the endocrine cells of the adrenal cortex, it also found that that Plin2-knockout adrenals had, an increased abundance of highly autofluorescent “ceroid like structures” in the inner region of the adrenal cortex containing lipid droplets, coupled with increased intra-adrenal cholesterol relative to wild-type animals. The present study indicates that these autofluorescent lipid-containing cell bodies are in fact cholesterol-containing BALL macrophages, a finding consistent with what is known about cholesterol containing-macrophages in other settings. Indeed, the degree of cholesterol esterification is known to increase autofluorescence in macrophages in the context of LDL uptake ^45^. Additionally, cholesterol-containing foamy macrophages in the context of atherosclerosis ^46^ are known to be autofluorescent, a feature that has been used to detect the cells in situ in the context of disease ^47^. Importantly, the Plin2 study ^20^ did not identify which cells are the source of Plin2 intra-adrenally. Our transcriptomic data indicate that Plin2 is expressed by adrenal macrophages, and therefore indicates that the autofluorescent cell phenotype observed in Plin2 knockout mouse adrenals was the primary phenotype, and not secondary to endocrine cells, as was presented in the study. Collectively these findings indicate that adrenal macrophage Plin2 is an additional molecular player in the regulation of BALL macrophage formation and adrenal hormonal output. This would appear to be a separate and distinct phenotype from that previously observed in relation to Liver X receptors. Liver X receptor α, which immunostaining indicates is expressed across endocrine cells in the adrenal cortex, is a negative regulator of adrenal cholesterol stores and corticosterone synthesis ^19^. Considering these findings collectively, it would suggest that BALL macrophages constitute a sizeable and functionally significant intra-adrenal cholesterol store that is distinct from, and additional to, the endocrine cell compartment’s own cholesterol stores. We have shown that this store contributes to hormonal output under steady state and in obesity, but the other contexts in which the store contributes to hormonal output remains to be explored.

Distinct macrophage subtypes, mostly CCR2-dependent monocyte-derived macrophages, have been previously linked with obesity-associated adipose tissue inflammation, insulin resistance, and hepatic steatosis ^29,48–50^. In the present study, our data suggest that CX3CR1-dependent BALL macrophages are responsible for the increase in corticosterone that comes with obesity ^51,52^. The data suggest therefore that BALL macrophages could represent a therapeutic target that could decouple obesity from the immune and metabolic dysfunction that is frequently synonymous with the condition. We have shown that these cells derive from a CX3CR1-expressing precursor, but whether this precursor is a yolk-sac erythro-myeloid progenitor, fetal liver progenitor, or haematopoietic stem cell-derived monocyte is currently unclear. Answering this question may be critical to inform attempts to target these cells for therapeutic benefit.

Obese mouse adrenals have previously been characterised. It was shown that adrenals from mice fed a high fat diet were significantly heavier, had cortical hyperplasia, and produced more corticosterone and aldosterone *ex vivo* than adrenals from mice fed a normal diet ^53^. However, this study did not control for the size of the adrenal gland. In our study, we report that adrenals from high fat-diet-fed CX3CR1^GFP/+^ and CX3CR1^GFP/GFP^ mice weigh the same yet differ in their corticosterone output that is measured in the plasma. This would therefore indicate that is the adrenal cholesterol content, but not overall size, that determines corticosterone output in the context of obesity.

A recent study also sought to characterise the adrenal macrophage compartment in depth ^37^. This study focused primarily on the sex-specific diversity of the macrophage compartment, but also established a link between macrophages and lipid metabolism. In contrast to the present study, Dolfi and colleagues ^37^ reported that global macrophage depletion reduced intra-adrenal, but not circulating, levels of aldosterone, and reported no effect on corticosterone. Notably, different experimental protocols were used in these studies, which can help explain the discrepancies in findings. The Dolfi study used a pan-macrophage targeting approach – anti-CD115 antibodies – to deplete adrenal macrophages in 8-week-old female mice and measured corticosteroid responses under cold challenge. We used a CX3CR1-targeting inducible diphtheria toxin receptor model to deplete ZGMs specifically in 10-week-old male mice, and measured corticosteroid responses at steady state under no challenge. This 8-10-week timepoint would also not be appropriate to assess the contribution of BALL macrophages to adrenal hormonal output as the timepoint is before significant accumulation of these cells (20 weeks +). Our study is therefore compatible with the Dolfi study and provides further mechanistic insight into how adrenal macrophages regulate corticosteroid output – via biosynthetic enzyme expression (ZGMs) and storage of the necessary corticosteroid precursor, cholesterol (BALLs).

While these findings constitute another intriguing example of the tissue-specific functions of macrophages, important questions remain outstanding. The precise mechanisms that regulate cholesterol accumulation in BALL macrophages remain unclear, as is the source of this cholesterol. We report that high fat diet feeding increases BALL size and cholesterol contents. This could be consistent with BALL macrophages scavenging cholesterol from the circulation via receptor-mediated mechanisms. Alternatively, adrenal macrophages could synthesise their own cholesterol, though this seems less likely, as they express key enzymes involved in cholesterol synthesis at very low levels. BALL macrophages could also recycle cholesterol from centripetally migrating endocrine cells ^54^, in a similar manner to the transient liver macrophages that recycle iron from senescent erythrocytes ^31^. Ultimately, further experimentation will be needed to establish the precise source of cholesterol that accumulates in BALL macrophages.

This study’s novelty lies in its demonstration of macrophages as their own distinct niche, with heterogeneity in form and function. This niche is highly specialised to the adrenal gland and serves to support adrenal gland hormonal output and could therefore introduce a new therapeutic paradigm in treating disorders driven by excessive corticosteroid responses. Additionally, this study provides yet another example illustrating that macrophages are more than “just” immune cells. Here, we present adrenal gland macrophages as additional players in the renin-angiotensin-aldosterone system and the hypothalamic-adrenal-pituitary axis. Whether or not macrophages in other endocrine organs serve similar functions remains to be explored.

## Acknowledgements

The authors would like to thank the University of Oxford Biomedical Services staff, The Don Mason Facility of Flow Cytometry (Sir William Dunn School of Pathology, University of Oxford), and the Light Microscopy Facility, Micron Advanced Bioimaging Unit (Dunn School, University of Oxford) for their help on this project. This research was funded by grants from Wellcome/HHMI International Research Scholar award 208576/Z/17/Z, ERC Consolidator grant – SympatimmunObesity ERC-2017 COG 771431, awarded to A.I.D, and the BHF graduate studentship FS/19/61/34900 awarded to C.J.O.O’B. For the purpose of Open Access, the author has applied a CC BY public copyright licence to any Author Accepted Manuscript (AAM) version arising from this submission.

## Author contributions

A.I.D conceptualised this study. C.J.O.O’B performed sample preparation, flow cytometry, body weight and adrenal weight measurements, and cholesterol and hormone measurements. D.O-S. performed single cell RNA-seq analysis. C.J.O.O’B. performed immunofluorescent confocal imaging and image analysis with the help of A.S., G.R, and C.A.P.S. P.L. and F.G. conceived of and carried out lineage tracing experiments. M.M., E.H.A., and L.R.S. carried out bulk RNA sequencing and data analysis. E.H. carried out RT-PCR. Y.M., T.I., and T.H. generated the Star-KO mouse adrenals and provided the tissue for analysis. E.P.G-S. and C.E.G-S. generated and provided the anti-CYP11B2 antibody and provided an intellectual contribution. S.G. provided mice and an intellectual contribution. D.R.G. provided mice, technical expertise and advice for the irradiation and bone marrow transplantation experiment. C.J.O.O’B. wrote the first draft of the manuscript and worked with A.I.D. to prepare the final version.

## Materials and methods

### Animals

C57BL/6J, CX3CR1-GFP, B6SJL CD45.1, and hCD68GFP/GFP mice were bred and housed in individually ventilated cages in specific pathogen-free conditions and on a a 12h light/dark cycle at 21°C+/-1°C, 50% humidity +/-10%. at the University of Oxford. All experiments on mice were conducted according to **institutional**, national, and European animal regulations. Animal protocols were approved by the animal welfare and ethics review board of the Department of Physiology, Anatomy, and Genetics at the University of Oxford. At the time of sacrifice, mice were anaesthetised with (10ul/g) 200mg/ml pentobarbitone sodium (Animalcare) solution, and cardiac perfused with 15-20mL of either 4% paraformaldehyde (PFA; used when adrenals were to be used for image analysis) or phosphate buffered saline (PBS; Sigma Aldrich; used when adrenals were processed for all other analyses, unless otherwise stated). All tissues were collected between 0930 and 1130. Prior to perfusion blood samples were taken from the right atria into EDTA-coated tubes. Blood samples were centrifuged at 1000g for 15 minutes to separate the plasma. Plasma samples were frozen at - 80°C until use in subsequent assays. Adrenal glands in all instances were dissected and any peri-adrenal adipose tissue was removed using a stereomicroscope. Adrenals to be used for microscopy and image analysis were fixed additionally overnight in 200uL 4% PFA at 4°C. After overnight fixation, PFA was removed from adrenals and adrenals were stored until use at 4°C in PBS containing 0.1% NaN_3_. Adrenals that were used in other experiments were frozen at −80°C until further analyses.

### FACS

Mice were euthanised by rising concentrations of CO_2_ and adrenals were harvested and any excess fat was trimmed. The cortex and medulla of dissected adrenal glands were separated manually using fine tweezers and scissors under a stereomicroscope. Cortices were minced using scissors and digested in 2.5 mg/mL type I collagenase (Sigma, C2674) and DNAse I for 1hr in a shaking 37oC incubator. Cells were live/dead stained using Fixable Aqua Live/Dead (Life Technologies, L34966), and non-specific antibody staining was blocked using Fc block (BD). Cells were surface stained using antibodies against CD45 (30-F11, BD), F4/80 (BM8, BioLegend), MHCII (M5/114.15.12, Invitrogen), CX3CR1 (SA011F11, BioLegend). Cells were permeabilised for intracellular staining using Foxp3 / Transcription Factor Staining Buffer Set (eBioscience) according to manufacturer’s instructions. Following permeabilization, cells were intracellular stained using antibodies against CD68 (FA-11, BD Biosciences) and Ki-67 (B56, BD Biosciences). Cell counts were determined using CountBright™ Absolute Counting Beads (Life Technologies) according to manufacturer’s instructions. UltraComp eBeads™ Compensation Beads (Invitrogen) were used to calculate compensation for flow cytometry. Samples were run on a BD Fortessa X20 flow cytometer. Flow cytometry data was analysed using FlowJo and Graphpad Prism 9 software.

### Cryosectioning and staining

Prior to tissue embedding, PFA-fixed adrenal glands were cryoprotected in 30% sucrose solution (in PBS) overnight. Cryoprotected adrenal glands were embedded in OCT embedding medium and snap frozen in liquid nitrogen. 15um sections were cut from embedded tissues using a Leica cryostat and mounted onto charged microscope slides. Cryosectioned tissue slices were thawed in PBS for 10 min at room temperature, tissue sections were circled using a super PAP pen (Life Technologies), blocked and permeabilised for 1 hour at room temperature in perm/block solution (3% bovine serum albumin, 2% goat serum, 1% Triton X-100, 0.01% NaN_3_ in PBS). Slides were stained with CD68-AF647 (clone FA-11, BioLegend), chicken anti-GFP (ab13970, Abcam) with goat anti-chicken AF488 (A11039, Invitrogen), and rabbit anti-CYP11B2 (rAS-2084, from Prof. Celso Gomez-Sanchez, University of Mississippi). Primary stains were done overnight at 4°C and secondary stains were done for 1 hour at room temperature. DAPI counterstains were done at 1:1000 for 5 minutes at room temperature. Fluoromount G mounting medium was used to mount coverslips to slides which were then sealed with clear nail varnish.

### Paraffin embedding, antigen retrieval, and antibody staining

Adrenal glands were first dehydrated in 70% (60 mins), 90% (45 mins) and absolute ethanol (45 mins), cleared with xylene (3x 60 mins), and infiltrated with Cellwax Plus paraffin wax (CellPath) at 60°C (2x 30 mins, 1x overnight). Samples were then embedded in cassettes in molten paraffin wax and left to cool. 10um sections were cut using a microtome, water mounted to charged slides, and melted onto slides for 1 hour. For antigen retrieval, first boil antigen retrieval solution (Stock: EDTA 10 mM SDS 0.5% pH 9.0, dilute 1:10 with Milli-Q water) with 900 Watt, afterwards place slides on a solution of EDTA 1mM, pH 9.0, SDS 0.05%, then heat up the solution again until it is boiling then sub-boil the buffer for 30 min 150 Watt in microwave. Slides were immediately washed and incubated in tap water for 5min each time, with regular gentle agitation. Slides were rinsed in distilled water prior to staining. Sections were blocked for 1 hour at room temperature with the blocking buffer as described above, and stained with rat anti-mouse CD68 antibody (clone FA-11, BioLegend) overnight at 4°C. Slides were washed in PBS and incubated with goat anti-rat alkaline phosphatase (Invitrogen, A18868) for 1 hour at room temperature. Fast Red staining was done using Liquid Fast-Red Substrate Kit (Abcam, ab64254) according to manufacturer’s instructions. Sections were counterstained with Mayer’s hematoxylin solution diluted 1:5 in water for 1 minute and washed in PBS. Fluoromount G mounting medium was used to mount coverslips to slides which were then sealed with clear nail varnish.

### Imaging and analysis

Slides prepared for immunofluorescence were imaged using a Zeiss 880 confocal microscope. IHC prepared slides were imaged using the EVOS M7000 imaging system. All image analysis was done using FIJI software on MacOS. Cell counting was done manually, with adrenal regions determined by differing cell morphologies. Quantitation of the CD68+ cell area and autofluorescence (raw integrated densities; RID) of immunofluorescent stains was done using FIJI image analysis software. 3-6 adrenal cryosections of stained tissue were analysed per mouse. CD68+ cells were segmented using the Otsu thresholding plugin and autofluorescent RIDs were taken from these segmented cells. Otsu was selected as it resulted in preferential and accurate segmentation of CD68++ macrophages. Otsu did not recognise smaller areas of CD68+ staining and so did not segment the dendritic-like zona glomerulosa macrophages, and so this method allowed for accurate, robust, unbiased, and reproducible cell size measurements. Cell size and autofluorescence from all slices were plotted as mean +/− s.e.m.

### Single cell sequencing analysis

Single-cell RNA-seq data from Dolfi *et al.* ^37^ were obtained from Gene Expression Omnibus (accession <u>GSE203095).</u> FASTQ sequencing libraries were used to generate digital gene expression matrices with 10x Genomics Cell Ranger 3.1.0, and the *scanpy* (PMID 29409532) python toolkit was used to orchestrate the analysis. For each sample (4 males, 4 females), cells were then filtering according to current quality control standards ^55^, and cells with at least 300 and less than 2,000 detected genes, less than 4,000 total mRNA counts and less than 5% of mitochondrial reads were kept for downstream analyses. Data for each cell were then library-size normalized and multiplied by a 10,000 scaling factor before log-normalization. Highly-variable genes were selected based on their dispersion and mean expression, and 3,166 genes were kept for embedding and clustering. Myeloid cells were detected by the expression of *Cd68, Cx3cr1, Ccr2 or Lyz2* and by the automatic cell-type assignment algorithm CellTypist^38^. After filtering myeloid clusters and subclustering, we identified an artefactual cluster that was visibly separate from the three others in the embedding and which was marked by the expression of *Pf4*, a platelet marker. After removal of this cluster, 6,377 high-quality myeloid cells were retained, comprising three separate clusters with specific marker genes. Embedding to a two-dimensional space was performed using Diffusion Maps (Ref Diffusion Maps). Gene expression was shown either on top of the embedding or as dotplots of standardized gene expression. For RNA velocity, we used Velocyto to quantify spliced and unspliced reads, and CellRank with standard parameters for the remaining analyses, the only particularity being the use of the Diffusion Components (DCs) as a latent orthogonal basis instead of Principal Component Analysis (PCA).

### Irradiation and bone marrow transplantation

10 week old male B6SJL CD45.1+ mice were irradiated with a total dose of 11Gy, which was administered as 2 doses of 5.5Gy, separated by 4 hours. These mice were injected intravenously with 9×10^6^ bone marrow cells from male hCD68-GFP mice ∼20 hours later. Mice were culled by CO_2_ 11 weeks later (at 20 weeks of age) and adrenals were harvested as previously described.

### Lineage tracing

For temporal lineage tracing experiments, 4-5 month old male Cx3cr1-CreERT2; Rosa26-mTmG mice were pulsed every 2 days for a total of five pulses to label Cx3cr1-expressing cells. Mice were aged for two further months prior to tissue harvest.

### FACS sorting for Bulk RNA sequencing

Male 22-week-old CX3CR1^GFP/GFP^ mice were euthanised and tissues harvested and processed for FACS as previously described. Endocrine (CD45-) and macrophage (CD45+ F4/80+ MHCII+ CX3CR1-GFP+) cell populations were sorted into Buffer RLT Plus (Qiagen) and RNA was extracted using the RNeasy Plus Micro Kit (Qiagen) according to manufacturer’s instruction.

### Library preparation, bulk RNA-seq mapping, gene counting and quality control

Reverse transcription, cDNA pre-amplification and sequencing library preparation were performed as previously described in [PMID: 35320714]. Briefly, reverse transcription and cDNA pre-amplification were performed using the SMART-Seq v4 Ultra Low Input RNA Kit for Sequencing (Clontech/Takara). cDNA was harvested and quantified with the Bioanalyzer DNA High-Sensitivity kit (Agilent Technologies). Libraries were prepared using the Nextera XT DNA Sample Preparation Kit and the Nextera Index Kit (Illumina) as per manufacturer instructions. Multiplexed libraries were pooled and paired-end 150-bp sequencing was performed on the Illumina HiSeq 4000 platform at Sidra Medicine

A standard RNA-Seq pipeline was launched using bcbio nextgen toolkit (https://doi.org/10.5281/zenodo.5781867). Briefly, sample Fastq files were mapped to the mouse reference genome GRCm38.p6 (GCA_000001635.8) using STAR (version 2.6.1d). Reads were retained only if uniquely mapped to the genome. FeatureCounts (v2.0.0) was used to obtain the number of reads that mapped to each gene. Sample Transcriptome data quality was assessed by evaluating homogeneity and similarity between samples. We used variance stabilizing transformation (vst) and regularized log transformation(rlog) functions (DESeq2 package (v1.26.0) ^56^) to transform the raw count data, (dist) function to calculate sample-to - sample distances. We plotted heatmaps of distance matrix along with PCA of 1000 most highly variable genes to identify putative outlier samples. These samples if present were excluded from the following differential expression analyses. We also used DESeq2 (v1.26.0) to normalize the raw count matrix according to sequencing depth and RNA composition (median of ratios method).

### Diphtheria toxin receptor-mediated macrophage ablation

Homozygous CX3CR1-Cre mice (Jackson Laboratories) were crossed with homozygous Gt(ROSA)26Sortm1(Dtr)ThBu (iDTR; R26iDTR) mice to generate heterozygous mice which express the diphtheria toxin receptor (DTR) from CX3CR1-expressing cells. At 10 weeks of age, 750ng Corynebacterium diphtheriaediptheria toxin (Calbiochem) or vehicle control was injected intraperitoneally in 200uL PBS. 24 hours later mice, were euthanised by pentobarbitone sodium solution as previously described, and blood samples were taken from the heart and processed as described earlier. Adrenals were harvested and processed for flow cytometry as previously described.

### Plasma hormone measurements

Plasma aldosterone and corticosterone concentrations were measured from plasma samples using the reverse competition ELISA kits (Enzo Life Sciences). Plasma norepinephrine was measured by ELISA (ImmuSmol). Plasma renin was measured using the Mouse Renin 1 (REN1) ELISA Kit (Invitrogen) and plasma ACTH was measured using Mouse/Rat ACTH SimpleStep ELISA® Kit (Abcam). All ELISAs were carried out according to the manufacturer’s instructions. Colorimetric and fluorimetric readouts from these assays were attained using a FLUOstar Omega plate reader (BMG Labtech).

### Cholesterol measurements

Plasma cholesterol was measured using the Cholesterol Quantitation Kit (Sigma Aldrich) according to manufacturer’s instructions. To prepare adrenal glands for cholesterol quantitation, whole adrenals were homogenised into 200ul chloroform:isopropanol:IGEPAL CA-630 (7:11:0.1) in a Precellys homogenisation tube (MK28-R). Tissues were homogenised using the PreCellys 24 tissue homogeniser. Homogenates were mixed thoroughly with 40ul sterile H2O, centrifuged at 13,000g for 10 minutes, and the upper aqueous phase was manually removed. The organic phase was transferred to a clean tube and vacuum dried at room temperature using a SpeedVac machine. Dried lipid pellets were dissolved with Cholesterol Assay Buffer and vortexed until homogenisation. The homogenised solution was used in the Cholesterol Quantitation Kit (Sigma Aldrich) according to manufacturer’s instructions.

### Temporal deletion of Star

Cre-ER^TM^; Star^flox/flox^mice were generated as previously ^41^. To induce knockout of Star in these mice, 8-week-old male Cre-ER^TM^; Star^flox/flox^mice were injected with 75 mg/kg body weight tamoxifen (Sigma Aldrich) three times every other day, then aged for 8 weeks prior to sacrifice.

### High fat diet challenge

At eight weeks old, mice of the indicated genotypes were given *ad libitum* access to a high fat diet (60% fat, D12492: Research Diets) for a period of 12 weeks. Tissues were harvested as described earlier for subsequent analyses.

### High salt diet challenge

For the high salt diet challenge experiment, 21-week-old CX3CR1^GFP/+^ mice were fed a diet containing 0.1% NaCl (diet D11112201; Research Diets) or a diet containing 2.5% NaCl (diet D22033001 with added NaCl; Research Diets) for six days. Tissues were harvested as described earlier for subsequent analyses.

### Subcutaneous osmotic minipump implantation

Osmotic minipump implantation was carried out as previously described ^42^. In short, Alzet osmotic mini-pumps (model 1002) containing vehicle or Ile5–Ang II (1000 ng/kg per minute, 14 days subcutaneous, Sigma Aldrich) were implanted into 25-week-old CX3CR1^GFP/+^ mice under isoflurane anaesthetisation. 14 days later mice were euthanised and adrenals and blood samples were harvested as described earlier.

### RT-qPCR

Adrenals were homogenised using the Precellys Hard Tissue Grinding Kit (MK28-R; Bertin Technologies). Total RNA from homogenised adrenals or lungs was isolated using RNeasy Plus Micro Kit (Qiagen, cat# 74034). cDNA was reverse transcribed using SuperScript II (Invitrogen) and random primers (Invitrogen). Quantitative PCR was performed using SYBR Green (Applied Biosystems) in C1000 TouchTM Thermal Cycler (BioRad). Β-actin was used as the housekeeping gene to normalize samples. We used the following formula to calculate the relative expression levels: RQ = 2^−ΔCt × 100 = 2−(Ct gene of interest – Ct β actin) × 100. The following mouse-specific primers were used. Star forward 5’-CAGGGCCAAGAAAACCTACA-3’; Star reverse 5’-ACGAGCATTTTGAAGCACCT-3’; Cyp11a1 forward 5’-AGGACTTTCCCTGCGCT −3’; Cyp11a1 reverse 5’-GCATCTCGGTAATGTTGG-3’; Hsd3b1 forward 5’-GCGGCTGCTGCACAGGAATAAAG-3’; Hsd3b1 reverse 5’-TCACCAGGCAGCTCCATCCA-3’; Cyp11b1 forward 5’-GGAAGAGAAGAGAGGGCAATGTGT-3’; Cyp11b1 reverse 5’-GGAAGAGAAGAGAGGGCAATGTGT-3’; Cyp11b2 forward 5’-CAGGGCCAAGAAAACCTACA-3’; Cyp11b2 reverse 5’-ACGAGCATTTTGAAGCACCT-3’; B-actin forward 5’-TCATGAAGTGTACGTGGACATCC-3’; B-actin reverse 5’-CCTAGAAGCATTTGCGGTGGACGATG-3’.

### LDL uptake assay

Bone marrow derived macrophages were generated as described previously ^57,58^. Briefly, bone marrow was obtained from tibae and femurs of 8-10 week old male C57Bl/6 mice or CX3CR1^GFP/GFP^ mice. Whole bone marrow isolates were cultured for 6 days in DMEM, 10% Fetal Bovine Serum (FBS), 10% L929 fibroblast-conditioned media as a source of M-CSF, 1% Penicillin/Streptomycin (P/S) in non-tissue culture treated 10cm petri dishes and incubated at 37°C, 5% CO_2_·. On day 5, 3 mL media was replaced with 5 mL fresh media. On day 6, cells were gently detached from dishes using a cell scraper, counted, and plated for experiments in DMEM containing 2% FBS in 96 well plates at a density of 100,000 cells per well. Cells were allowed to settle overnight before subsequent experimentation. On day 7, BMDM cultures were treated with 250ug/uL BODIPY™ FL complex Low Density Lipoprotein (Invitrogen) for 24 hours. Cells were then fixed with 4% PFA for 15 minutes, washed with PBS, and fluorescence intensity was measured using the FLUOstar Omega plate reader (BMG Labtech; Ex/Em 515/520).

### Statistical Analysis

All experiments, where possible, were designed to have equal groups of biological replicates (n) per experimental group. Data in all instances were presented as mean +/− standard error mean (s.e.m.). Technical replicates were used in plate-based assays, the average of which was used to represent each biological n. Statistical analysis was done using Graphpad Prism 10.0 for MacOS, GraphPad Software, Boston, Massachusetts USA, www.graphpad.com. For comparison of two experimental groups, an unpaired student’s t test was used. For comparison of more than two groups, a one-way ANOVA with Tukey’s post-hoc test for multiple comparisons. Significance in all cases was denoted as follows: \**P* < 0.05, \*\**P* < 0.01, \*\*\**P* < 0.001, \*\*\*\**P* < 0.0001.

---

**Supplementary figure 1.**
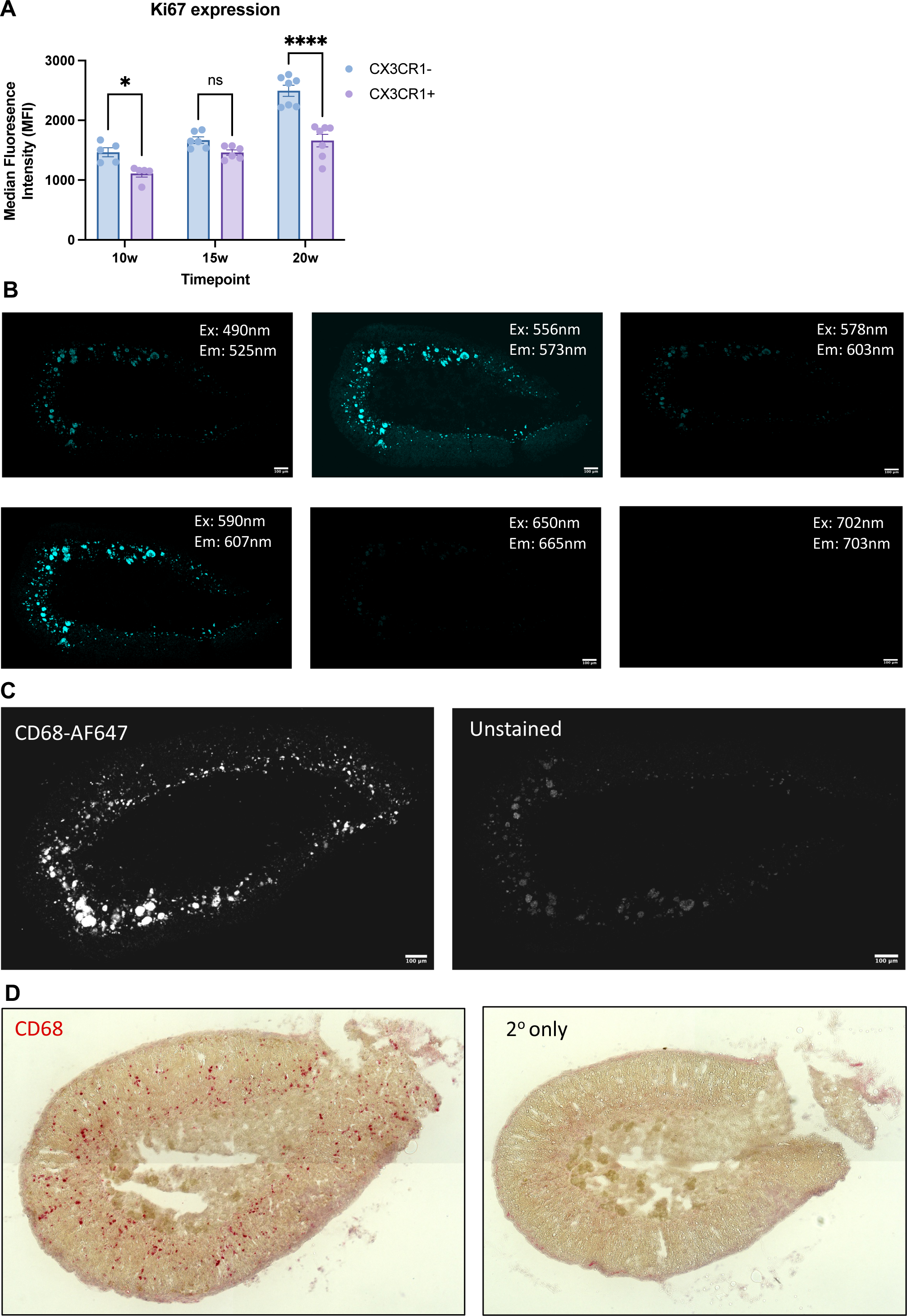
(A) Flow cytometric data showing the median fluorescence intensity of Ki-67 expression in the indicated macrophage subpopulations at the indicated timepoints. (B) Confocal images showing the autofluorescent signals at the indicated excitation/emissions from unstained 22-week-old male adrenal glands. (C) Confocal images showing a cross section of a 22-week-old male wild-type murine adrenal gland stained with anti-CD68-AF647 (white) (left) and an unstained slice (right) taken from the same mouse adrenal gland. (D) An optical micrograph of a 22-week-old male mouse adrenal gland stained with H&E and ant-CD68 Fast Red staining (left) and a secondary antibody-only control (right). Scale bars are 100μm. Data in A were analysed by one way ANOVA with Tukey’s test for multiple comparisons and are shown as average ± s.e.m. \**P* < 0.05, \*\**P* < 0.01, \*\*\**P* < 0.001, \*\*\*\**P* < 0.0001.

**Supplementary figure 2.**
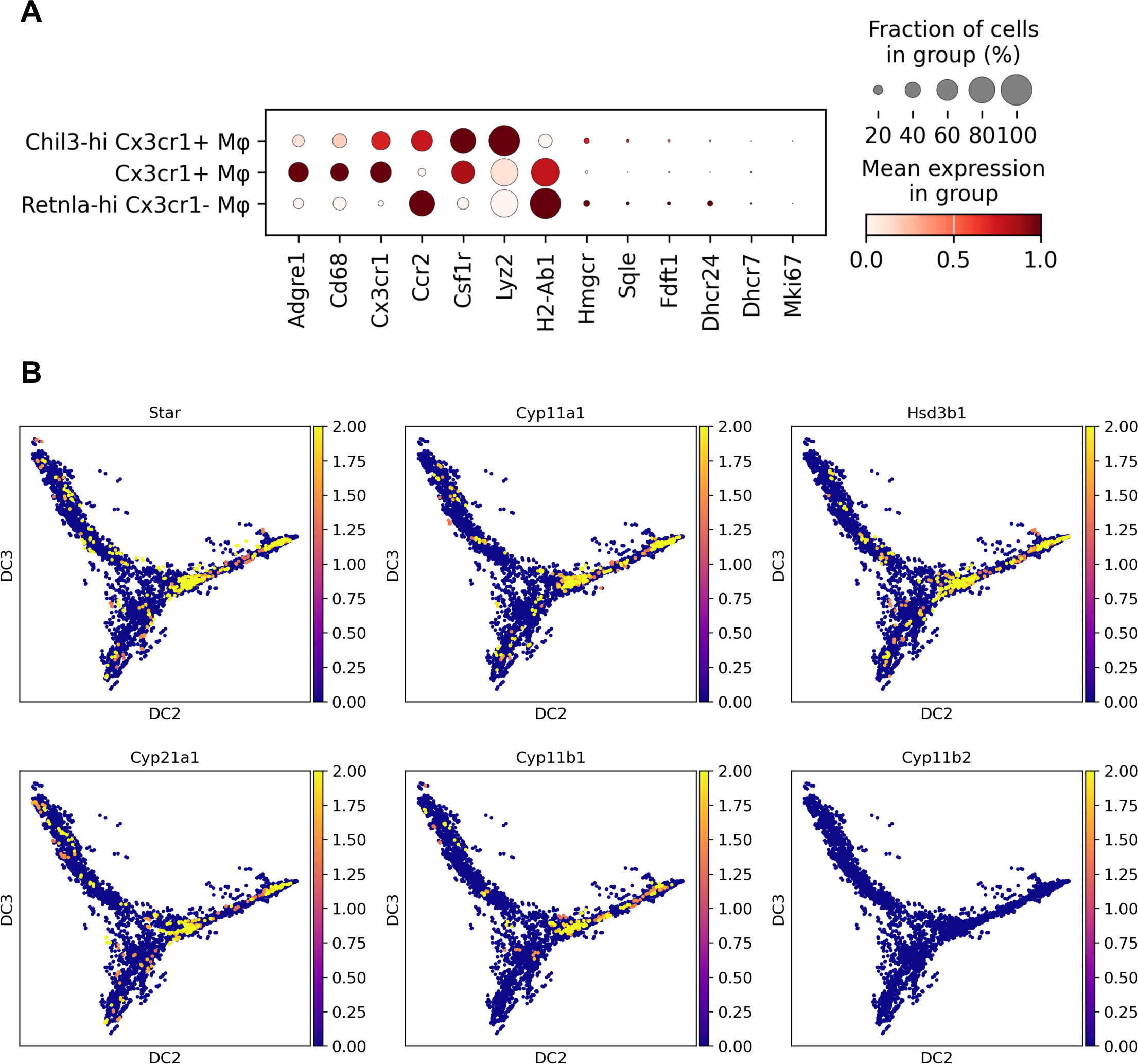
(A) ScRNAseq dotplot showing the standardised gene expression of macrophage marker genes and cholesterol biosynthetic genes in adrenal myeloid cells. (B) Gene expression data showing the expression of corticosteroid biosynthetic enzymes in adrenal myeloid cells.

**Supplementary figure 3.**
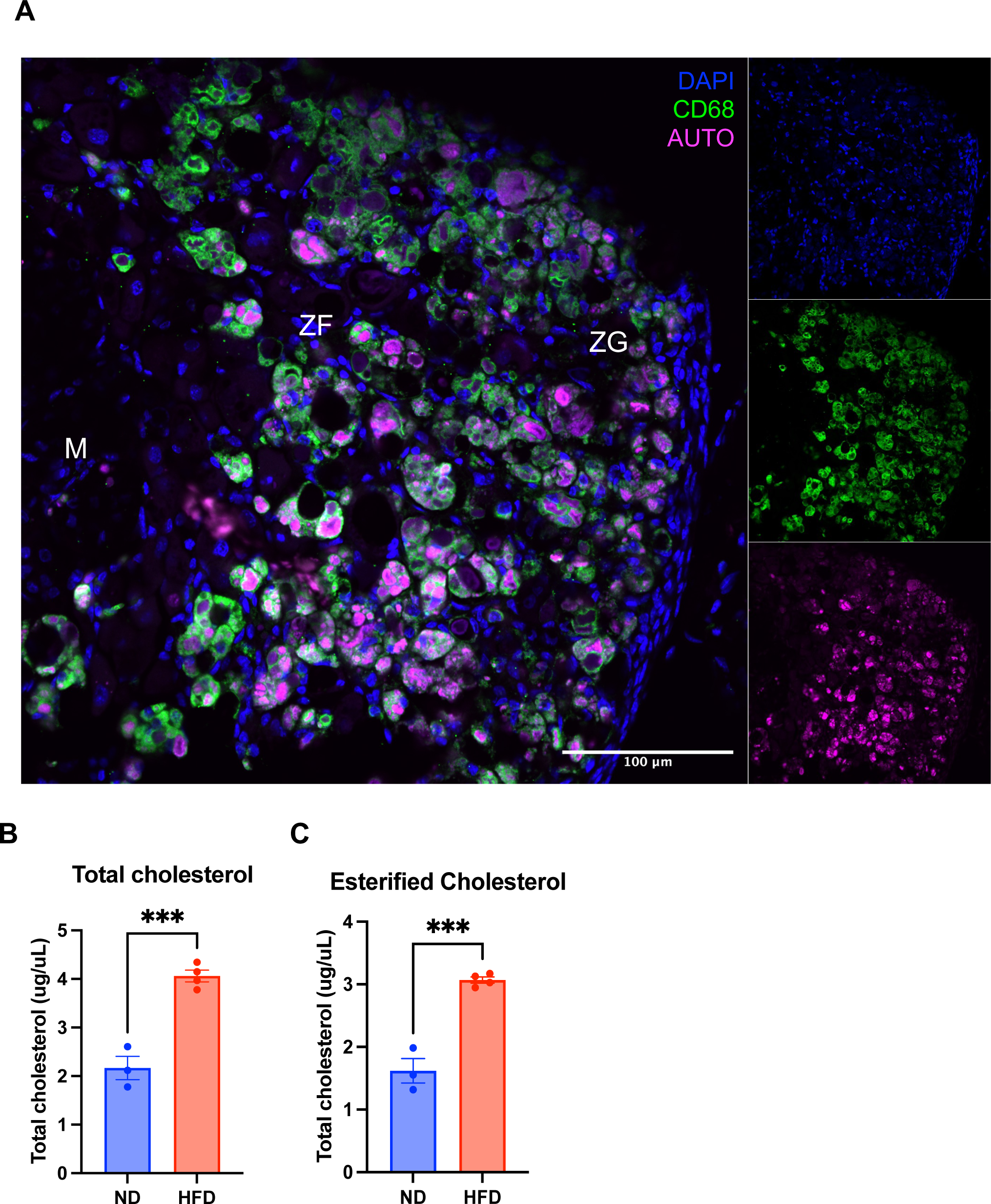
(A) A representative confocal image showing an adrenal gland from a CAGGCre-ER^TM^; Star^flox/flox^ mouse stained with anti-CD68 (green) and DAPI (blue), with autofluorescence (magenta). High fat diet increases total plasma cholesterol (B) and cholesterol esters (C). ZG = zona glomerulosa; ZF = zona fasciculata; M = medulla; AUTO = autofluorescence.

**Supplementary figure 4.**
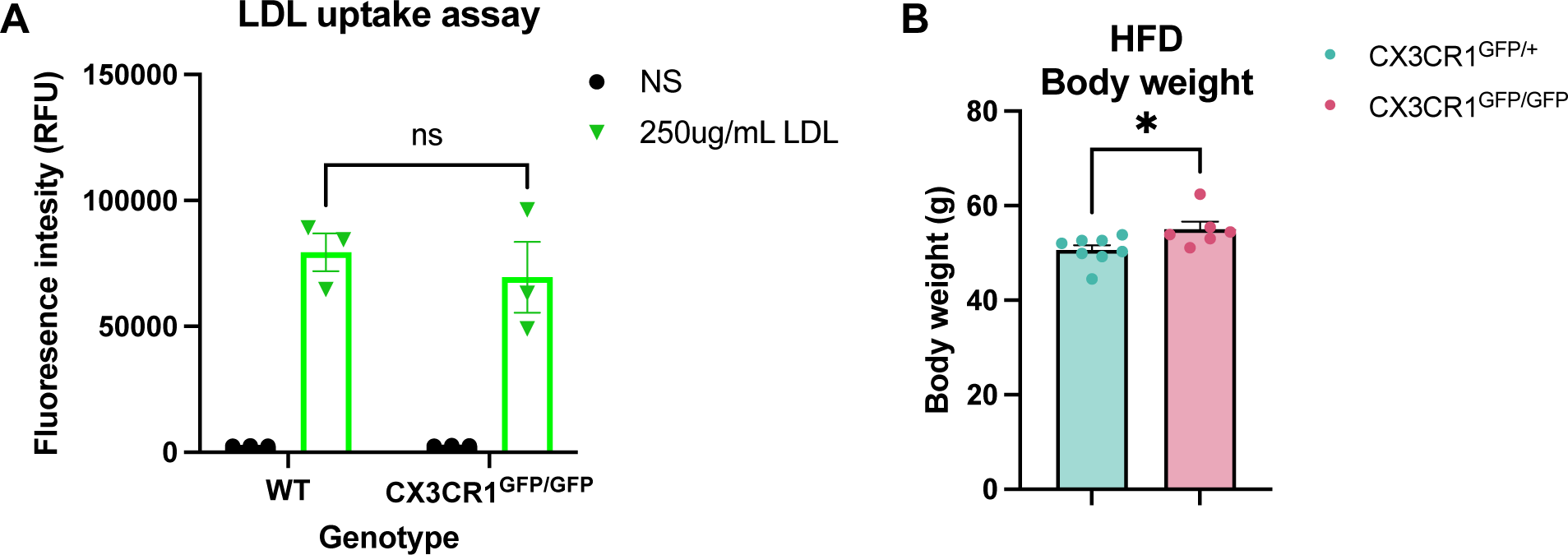

